## Supplementary Table S2 for "Eicosanoid content in fetal calf serum accounts for reproducibility challenges in cell culture"

**Supplementary Table S2:** Inclusion list of eicosanoids as used for the MS/MS analysis

Mass [m/z]

254,2245

275,2011

277,2167

279,2324

281,2480

283,2637

293,2122

295,2279

301,2168

303,2330

311,2228

315,1966

317,2122

319,2279

321,2435

327,2324

327,2781

329,2480

333,2071

335,2222

337,2384

343,2279

348,3069

349,202

351,2177

353,2328

355,2428

357,2585

359,2222

367,3576

375,2171
