## Supplementary Figure S1 for "Eicosanoid content in fetal calf serum accounts for reproducibility challenges in cell culture"

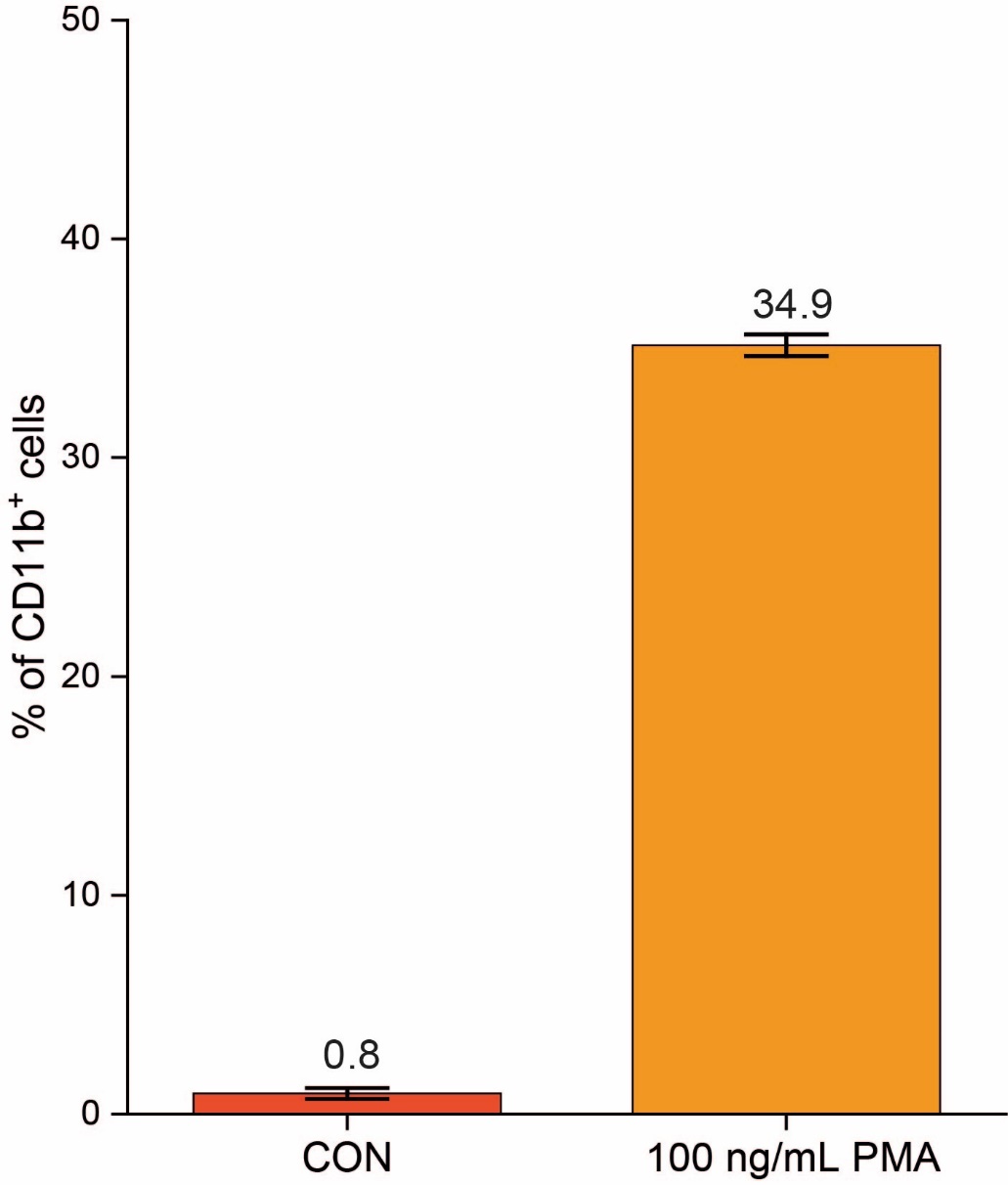

**Supplementary Figure S1:** Percentage of CD11b (ITGAM) positive U937 cells after 48 h treatment with 100 ng/mL PMA. The analysis shows a significant (p-value<10^-8^) increase in CD11b^+^ cells upon treatment compared to untreated cells.

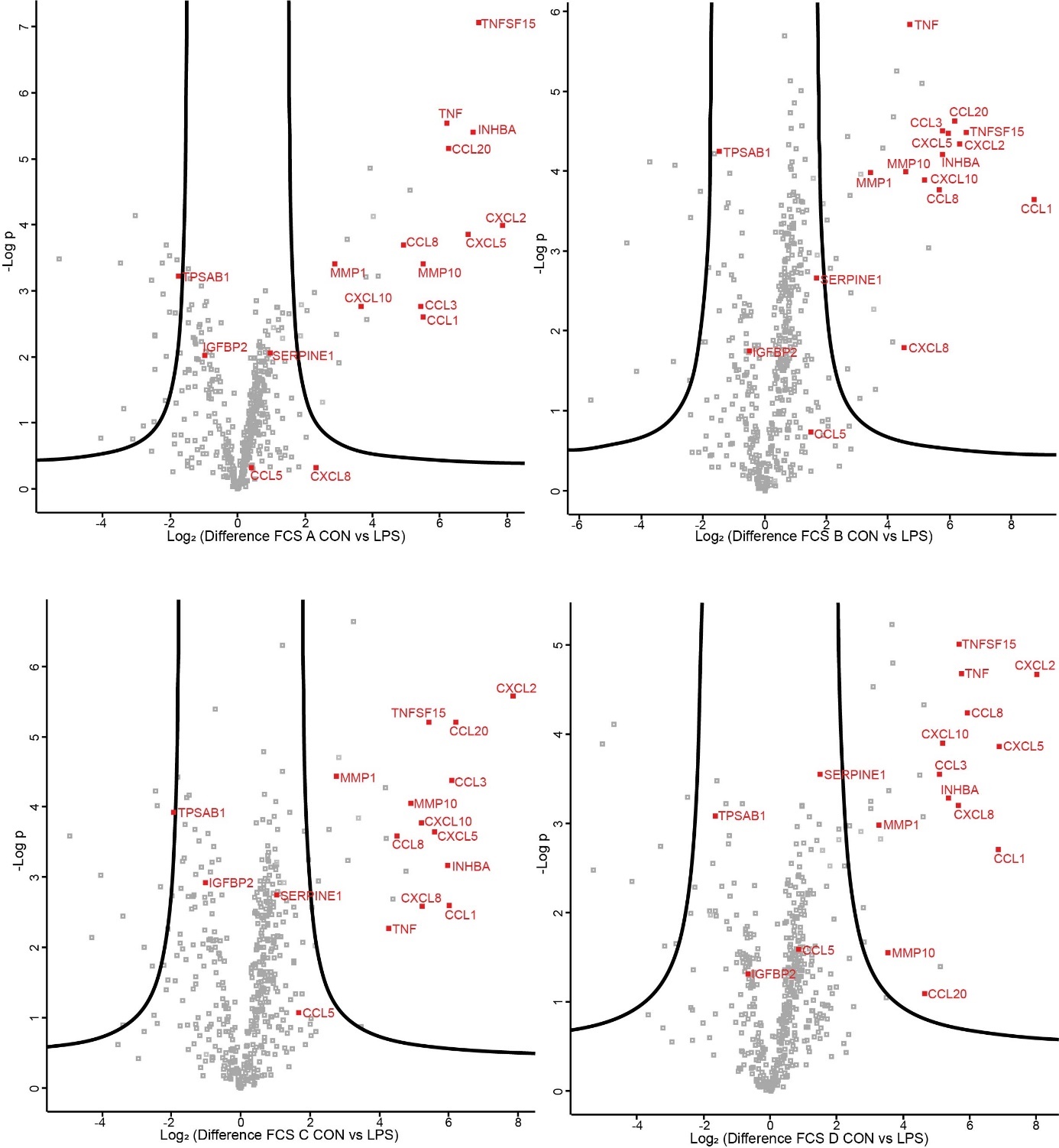

**Supplementary Figure S2:** Volcano plots comparing the identified proteins of the control and the LPS activated U937 cells for each medium A-D. The significant regulated proteins (FDR=0.05, S0=2) were used for the Venn diagram shown in Figure 1C. The labelled proteins are described in the context of inflammatory activation and are mentioned in the main text. All identified proteins and the significant regulations with respective p-value and differences can be found in supplementary table 3.

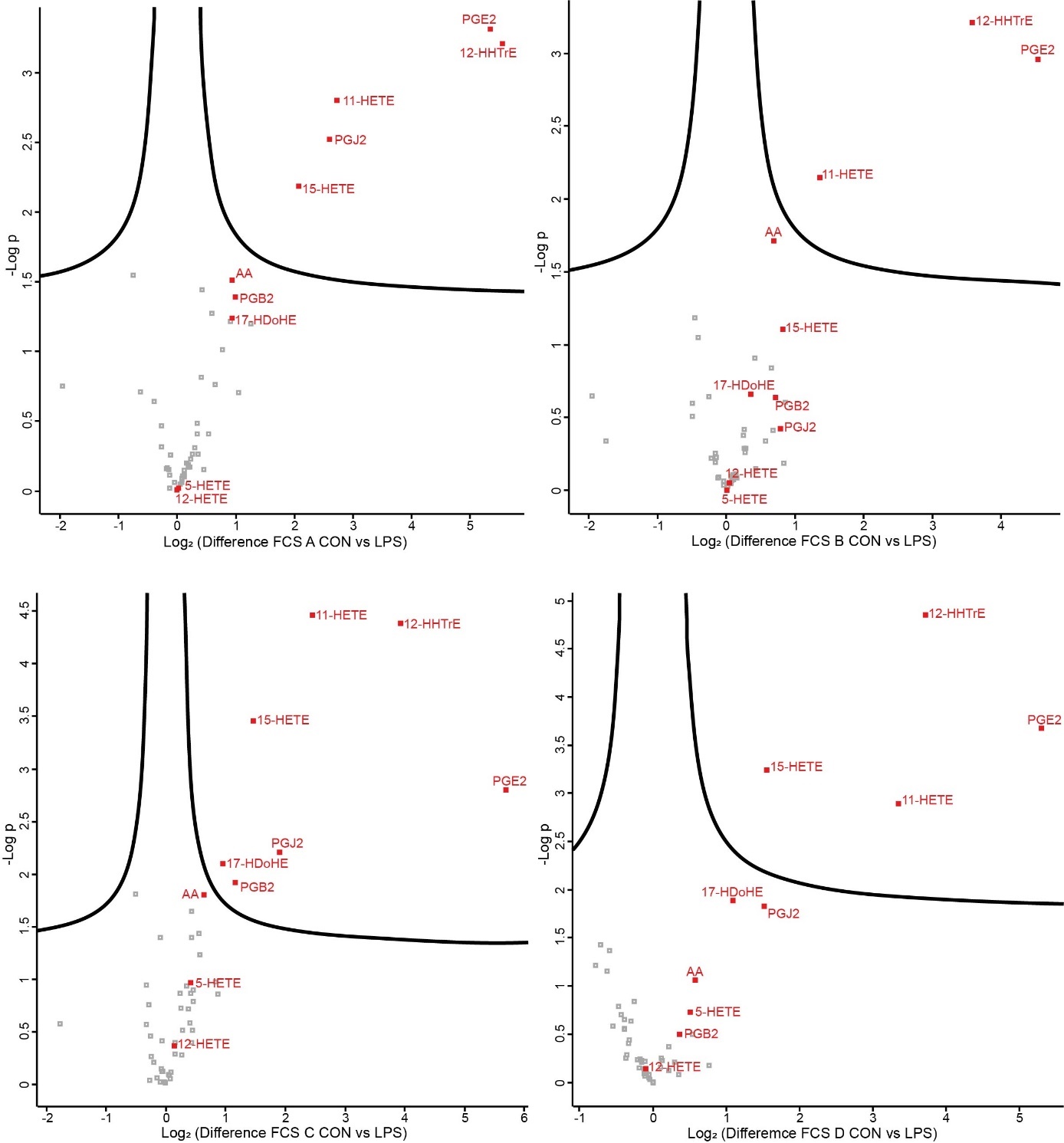

**Supplementary Figure S3:** Volcano plots comparing the identified eicosanoids of the control and the LPS activated U937 cells for each medium A-D. The significant regulated oxylipins (FDR=0.05, S0=0.1) were used for the Venn diagram shown in Figure 1D. All identified oxylipins and the significant regulations with respective p-value and differences can be found in supplementary table 4.
