## Supplementary Table S1 for "Eicosanoid content in fetal calf serum accounts for reproducibility challenges in cell culture"

**Supplementary Table S1:** Eicosanoid Standards; All eicosanoid standards were bought from Cayman Europe, Tallinn, Estonia.

| **Standard Mixture** | **Name** | **Abbreviation** | **Conc. Stock solution [nM]** | **Final conc. [nM]** |
| --- | --- | --- | --- | --- |
| Standards spiked **before** solid phase extraction | 12S-Hydroxyeicosatetraenoic acid d8 | 12S-HETE-d8 | 609,2 | 20,3 |
|  | 15S-Hydroxyeicosatetraenoic acid d8 | 15S-HETE-d8 | 609,2 | 20,3 |
|  | 5-Oxo-Eicosatetraenoic acid d7 | 5-Oxo-ETE-d7 | 1844,6 | 61,5 |
|  | 11,12-Dihydroxy-5,8,14-icosatrienoic acid d11 | 11.12-DiHETrE-d11 | 1122,8 | 37,4 |
|  | Prostaglandin E2-d4 | PGE2-d4 | 613,0 | 20,4 |
|  | 20-Hydroxyeicosatetraenoic acid d6 | 20-HETE-d6 | 572,5 | 19,1 |
| Standards spiked **after** solid phase extraction | 5S-Hydroxyeicosatetraenoic acid d8 | 5S-HETE-d8 | 609,2 | 20,3 |
|  | 14,15-Dihydroxy-5,8,11-icosatrienoic acid d11 | 14.15.DiHETrE-d11 | 572,5 | 19,1 |
|  | 8-Isoprostaglandin F2α d4 | 8-iso-PGF2α-d4 | 1116,5 | 37,2 |
